# Mutations on the surface of HDAC1 reveal molecular determinants of specific complex assembly and their requirement for gene regulation

**DOI:** 10.1101/2025.02.24.639909

**Authors:** Ahmad Alshehri, India-May Baker, David M English, Louise Fairall, Mark O Collins, John WR Schwabe, Shaun M Cowley

## Abstract

Histone deacetylase 1 and 2 (HDAC1/2) are highly related enzymes that regulate histone acetylation levels in all cells, as catalytic and structural components of six unique multiprotein complexes: SIN3, NuRD, CoREST, MIDAC, MIER and RERE. Co-immunoprecipitation of HDAC1-Flag followed by mass spectrometry revealed that 92% of HDAC1 in mouse embryonic stem cells resides in 3 complexes, NuRD (49%), CoREST (28%) and SIN3 (15%). We compared the structures of MTA1:HDAC1 and MIDAC:HDAC1 to identify critical binding residues on the surface of HDAC1. Surprisingly, a single mutation, Y48E, disrupts binding to all complexes except SIN3. Rescue experiments performed with HDAC1-Y48E in HDAC1/2 double-knockout cells, showed that retention of SIN3 binding alone is sufficient for cell viability. Gene expression and histone acetylation patterns were perturbed in both Y48E and a second mutant cell line, HDAC1-E63R, indicating that cells require a full repertoire of the HDAC1/2 complexes to regulate their transcriptome appropriately. Comparative analysis of MTA1/HDAC1 and SIN3B/HDAC2 structures confirmed the differential modes of HDAC1 recruitment, such that Y48 interacts with ELM2/SANT domain-containing proteins, but not SIN3. The E63R mutation shows markedly reduced binding to NuRD and MiDAC complexes, but retains some CoREST binding. We provide novel molecular insights into the abundance, co-factors and assemblies of this crucial family of chromatin modifying machines.

## Introduction

The post-translational modification of histones is a fundamental mechanism for the regulation of chromatin organisation in all eukaryotes. Acetylation of highly conserved lysines in the N-terminal tails of all four core histones leads to a loosening of chromatin structure as well as providing binding sites for acetyl-lysine reader proteins ^1^. Critically, histone acetylation is a dynamic process, with a half-life of 30 to 120 mins ^2,3^, due to the interplay of histone acetyltransferase (HAT) and histone deacetylase (HDAC) enzymes. HDAC1 and 2 (HDAC1/2) are highly-conserved paralogues that serve as common catalytic components of six distinct multiprotein complexes: SIN3, NuRD, CoREST, MiDAC, MIER and RERE. Each consists of a central platform (e.g., SIN3A) that mediates the binding of HDAC1/2 and numerous auxiliary proteins that add complementary activities ^4,5^. The majority of these complexes can be further divided into sub-complexes, often by switching interchangeable central platforms, SIN3A versus SIN3B for instance ^6^. This is not just a simple case of gene duplication and redundancy. SIN3A and SIN3B produce different mouse knockout phenotypes ^7–9^ and biochemically discrete complexes in cells, with a unique array of SIN3-associated proteins ^10^. An analogous arrangement also occurs in the NuRD and CoREST complexes, with interchangeable central components, MTA1-3 and RCOR1-3 respectively, and an assortment of supplementary factors, including enzymes such the helicase, CHD4 (NuRD) and lysine-specific demethylase 1 (LSD1/KDM1A) in CoREST.

The 15 HDAC1 binding partners can be split into two main classes, the majority that employ ELM2-SANT domains (MTA1-3, RCOR1-3, MIER1-3, MIDEAS, TRERF1, ZNF541 and RERE) and the two highly related corepressors, SIN3A and SIN3B. With the exception of ZNF541 (testis-specific), the expression of these proteins is largely ubiquitous in human cells. Each cell type, therefore, has the option to employ 14 different HDAC1-binding platforms, with a multitude of accessory proteins, contributing both chromatin binding and modifying activities. In mice, knockout of critical components of the SIN3A, NuRD, CoREST and MIDAC complexes results in embryonic lethality ^7,11–14^, suggesting that each possesses a unique and essential function. Why cells contain such a variety of vehicles for HDAC1/2 activity remains to be fully understood. Structural biology has provided insights into the assembly of these macromolecular machines. In addition to a unique set of components, complexes have either monomeric (CoREST), dimeric (SIN3A, NURD) or tetrameric (MiDAC) assemblies ^12,15–17^, particularly pertinent for nucleosomal substrates containing two copies of each core histone.

To disentangle unique from overlapping activities of HDAC1/2 complexes, we require additional information about complex components, their relative abundance and mode of assembly. For example, are all complex components present in the same amount? And if not, what constitutes a *core* complex? To address these questions, we have performed a quantitative mass spectrometry analysis of HDAC1/2 complexes purified from mouse embryonic stem cells and found that the majority of HDAC1 is present in just 3 complexes (NuRD, CoREST and SIN3A), with a relatively small set of unique core components. Their assembly utilises HDAC1 as an integral structural component, with conserved residues on the surface of the enzyme contributing to a highly stable association. Comparative structural analysis of individual ELM2-SANT domains bound to HDAC1 allowed the identification of key surface residues. Surprisingly, given an extensive protein-protein interaction surface, we found that single point mutations on the surface of HDAC1 are sufficient to perturb binding to its partners. Furthermore, by interrogating individual binding regions, we have identified the first mutations of their kind that allow us to discriminate between different HDAC1 binding proteins and their associated complexes. By probing how HDAC1/2 complexes are assembled, we have gained insights into their unique functions, and the first indications of how to perturb specific complexes inside cells.

## Methods

### Cell culture

All experiments described in this research used *Hdac1/2^lox/lox^* ES cells cultured as described previously in M15 + leukaemia inhibitory factor (LIF) media ^18^. These cells were engineered to stably express either wild-type HDAC1 rescue or HDAC1 mutants as indicated. To induce deletion of endogenous *Hdac1/2*, cells were cultured with 1 μM 4-hydroxytamoxifen (OHT) for 24 hours and then cultured for a further 4 days to ensure complete removal of endogenous HDAC1/2 before use in the experiments detailed below.

### Lipofectamine 2000 stable transfections

The cloning of all constructs used in this work was performed by the PROTEX service at the University of Leicester. Stable transfection was achieved by co-transfecting Hdac1/2*^lox/lox^* cells with *piggyBac* transposase and expression vectors as described previously ^19^.

### CellTiter-Glo (CTG) assays

For each HDAC1 mutant, 500 cells were seeded per well in triplicate on 96 well plates (−/+ OHT). Five days later CTG assays (Promega) were conducted as per the manufacturer’s instructions.

### Crystal Violet viability assays

For each HDAC1 mutant, 8 × 10^5^ cells were seeded into 6 well plates in triplicate (−/+ OHT). 5 days post OHT treatment media was removed and cells were washed once with PBS prior to fixing with 1 mL of 100% methanol for 2 mins. Methanol was removed and cells were washed once with PBS before being left to dry for 3 hours at room temperature. 1 mL Crystal Violet reagent (ProLab; PL.7000) was then added to each well for 2 mins then cells were washed with water. Plates were allowed to dry for 24 hours before imaging.

### Co-Immunoprecipitations

Five days prior to the coIPs, HDAC1 rescue cell lines were treated with 1 μM OHT to remove endogenous HDAC1/2. Cell pellets from a confluent 10 cm plate were lysed in an appropriate volume of coIP buffer (250 mM NaCl, 20 mM Tris-HCl pH 7.5, 0.5% IGEPAL, 1 mM EDTA, Protease inhibitor (Roche; 05892791001)) on ice for 30 mins. Lysed samples were then centrifuged at 14,000 rpm at 4°C for 15 mins, the supernatant containing the protein extract was retained and the concentration was determined by Bradford assay. Extracts were diluted to a concentration of 5 μg/μL using coIP buffer and 300 μL was used per IP. 60 μL of Protein G Dynabeads (Thermo Fisher; 10003D) was washed 3 times with coIP buffer and then resuspended in an equal volume of coIP buffer to the original volume of bead slurry before the addition of 1 μg of Flag or normal mouse IgG antibodies. Antibodies were bound to beads via rotation at 4°C for 1 hour. Beads were again washed in 500 μL of coIP buffer and then resuspended in 200 μL of coIP buffer before being split equally between IPs. IPs were incubated at 4°C overnight (18 hours) with rotation, 20% of the protein extract was kept as an input before the addition of beads. IPs were then washed 3 times with coIP buffer before being used for western blotting or mass spectrometry as described below.

### Western blotting

Western blots were performed as described previously ^18^. Where coIP samples were used for western blotting, the beads were resuspended in 100 μL of coIP buffer before the addition of 4x loading buffer (Invitrogen; NP0007) and denaturation at 95°C for 5 mins.

### Preparation of HDAC1-Flag pulldown samples for mass spectrometry analysis

HDAC1-Flag pulldown samples were eluted by heating in 5% SDS, 50 mM Tris pH 7.4 at 70°C for 20 minutes, Triethylammonium bicarbonate (TEAB) was then added to a final concentration of 50 mM with a pH of 8. Tris(2-carboxyethyl) phosphine hydrochloride (TCEP) was added to a final concentration of 5 mM and samples were heated to 70 °C for 15 minutes. Iodoacetamide (IAA) was then added to a final concentration of 10 mM, and samples were incubated at 37 °C in the dark for 30 minutes. Samples were acidified with 4 μL of 12 % phosphoric acid and 264 μL of S-Trap binding buffer (90% methanol, 100 mM TEAB pH 7.1) was then added to each sample. Samples were then passed through S-Trap columns 150 μL at a time by centrifugation at 4000 rpm for 30 seconds. Samples were washed four times with 150 μL of S-Trap binding buffer through centrifugation at 4000 rpm for 30 seconds. 2 μg of trypsin (Pierce, sequencing grade) was added to each sample, and digestion was allowed to proceed at 47 °C for 80 minutes. Peptides were eluted with 40 μL of 50 mM TEAB, 40 μL of 0.2% aqueous formic acid and 40 μL of 50% acetonitrile with 0.2% formic acid, with centrifugation at 4000 rpm for 30 seconds for each elution. Eluted peptides were dried in a vacuum concentrator and resuspended in 0.5% formic acid for LC-MS/MS analysis.

### Preparation of histone samples for mass spectrometry analysis

To obtain histone proteins a whole cell extract was first made by lysing cell pellets from confluent 10 cm plates in NP-40 lysis buffer (50 mM Tris-HCl pH 8, 150 mM NaCl, 1 mM EDTA, 1% NP-40, 10% glycerol, protease inhibitor cocktail (Sigma; P8340)) for 30 mins. Lysates were cleared by centrifugation at 14,000 rpm at 4 °C for 15 mins. The supernatant containing the whole cell extract was transferred to a fresh tube and histones were extracted from the remaining pellet by overnight incubation (20 hours) in 50 μL of 0.4N H_2_SO_4_. Samples were again centrifuged at 14,000 rpm at 4 °C for 15 mins, and the supernatant containing the histone proteins was transferred to a fresh tube. The samples were then neutralised using 0.8N NaOH. Histones were derivatised according to ^20^. Briefly, 10 μg of histone samples in 50 μL of 50 mM ammonium bicarbonate was incubated with 16.6 μL of propionylation reagent (1:3 v/v propionic anhydride in acetonitrile) for 15 mins at 37°C with shaking at 900 rpm. Samples were dried down in a vacuum concentrator and the derivatisation steps were repeated. 10 μg of derivatised histone samples were digested with 1 μg of trypsin in 50 μL of 50 mM ammonium bicarbonate for 2 hours at 37°C with shaking at 900 rpm. Digests were desalted using C18 spin columns (Thermo Fisher; 89870) according to the manufacturer’s protocol. Eluted peptides were dried in a vacuum concentrator and resuspended in 0.5% formic acid for LC-MS/MS analysis.

### Mass spectrometry analysis

Each sample was analysed using nanoflow LC-MS/MS using an Orbitrap Elite (Thermo Fisher) hybrid mass spectrometer equipped with an EasySpray source, coupled to an Ultimate RSLCnano LC System (Dionex). Peptides were desalted online using a nano-trap column, 75 μm I.D.X 20mm (Thermo Fisher) and then separated using a 120-min gradient from 5 to 35% buffer B (0.5% formic acid in 80% acetonitrile) on an EASY-Spray column, 50 cm × 50 μm ID, PepMap C18, 2 μm particles, 100 Å pore size (Thermo Fisher). The Orbitrap Elite was operated in DDA mode with a cycle of one MS (in the Orbitrap) acquired at a resolution of 120,000 at m/z 400, a scan range 375-1600, with the top 20 most abundant multiply charged (2+ and higher) ions in a given chromatographic window subjected to MS/MS fragmentation in the linear ion trap using CID. An FTMS target value of 1e6 and an ion trap MSn target value of 1e4 were used with the lock mass (445.120025) enabled. Maximum FTMS scan accumulation time of 200 ms and maximum ion trap MSn scan accumulation time of 50 ms were used. Dynamic exclusion was enabled with a repeat duration of 45 s with an exclusion list of 500 and an exclusion duration of 30 s.

### Mass spectrometry data analysis

Raw mass spectrometry data were analysed using MaxQuant version 1.6.10.43. The following parameters were used to search against a mouse reference proteome: digestion set to Trypsin/P with 3 missed cleavages, methionine oxidation, N-terminal protein acetylation and lysine acetylation set as the variable modifications. Additionally, propionylation was set as a variable modification for derivatised histone samples with the number of missed cleavages set to 5. A protein and peptide FDR of 0.01 were used for identification level cut-offs based on a decoy searching database strategy. Protein group output files generated by MaxQuant were loaded into Perseus version 1.6.10.50. The matrix was filtered to remove all proteins that were potential contaminants, only identified by site and reverse sequences. The LFQ intensities were then transformed by log2(x), normalised by subtraction of the median value, and individual intensity columns were grouped by experiment. Proteins were filtered to keep only those with a minimum of 3 valid values in at least one group. The distribution of intensities was checked to ensure standard distribution for each replicate. Missing values were randomly imputed with a width of 0.3 and downshift of 1.8 from the standard deviation. To identify significant differences between groups, two-sided Student’s t-tests were performed with a permutation-based FDR of 0.05. The mass spectrometry data have been deposited to the ProteomeXchange Consortium via the PRIDE partner repository with the dataset identifiers (PXD060154 (HDAC1 pulldowns) and PXD060158 (histone analysis))

### Deacetylase assays

In vitro HDAC assays were conducted on whole cell extracts made from cells expressing the different HDAC1 mutants extracted in HDAC assay buffer containing 50 mM NaCl, 50 mM Tris-HCL pH 7.5, 5% Glycerol, 0.3% Triton X-100 using a Boc-Lys(Ac)-AMC substrate as described previously ^18^. One-way ANOVA multiple hypothesis testing was performed using GraphPad Prism to determine significant changes in deacetylase activity (n=3).

### RNA-sequencing

RNA was extracted from cell pellets collected from a 6 cm plate using the TRI Reagent (Zymo Research, R2050-1-200) based Direct-Zol RNA MiniPrep Kit (Zymo Research, R2052) following the manufacturer’s instructions. RNA quality was determined by the NUCLEUS facility at the University of Leicester using an Agilent RNA 6000 Nano Chip on an Agilent 2100 Bioanalyzer. Library preparation and sequencing was carried out by Novogene, samples were sequenced to a depth of 20 million reads on the Illumina NovaSeq 6000 PE150 sequencing platform. Bioinformatic analysis was performed as described in ^21^ with reads mapped to the mm10 genome index. The processed files can be found at the Gene Expression Omnibus (GEO), series GSE278462. For gene ontology (GO) analysis, the Bioconductor package TopGO ^22^ was used with the mouse genome-wide annotation package org.Mm.eg.db.

## Results

### NuRD, CoREST and SIN3A are the dominant HDAC1/2 complexes in cells

To examine the composition and stoichiometry of HDAC1/2 containing complexes, we stably expressed HDAC1 with a C-terminal Flag tag in conditional *Hdac1/2* double knockout (DKO) embryonic stem cells (ESCs) (Fig 1A). Treatment with 4-hydroxytamoxifen (4OHT) activates removal of the endogenous HDAC1/2 proteins within 4 days ^18^, allowing us to assess the proteins bound to HDAC1-Flag using co-immunoprecipitation (coIP) and mass spectrometry (Fig 1A). We identified 148 proteins significantly enriched in the pulldown, compared to DKO cells lacking HDAC1-Flag as a control (supplementary Table S1). Components of all six HDAC1/2 complexes were present among this list. To estimate the abundance of individual proteins we calculated intensity Based Absolute Quantification (iBAQ ^23^) values and manually curated proteins into SIN3A/B, NuRD, CoREST, MiDAC, MIER and RERE complexes (Fig 1B and 1C). Of the 14 HDAC1 interacting proteins expressed in ESCs, we detected 13 (Fig 1B), with only RCOR3 absent, most probably due to lower expression levels. Using the iBAQ value for direct-HDAC1 binding proteins in each complex we estimated the relative abundance of the six complexes (Fig 1B, indicated by red arrows). Almost half of the HDAC1 in ESCs was incorporated into the NuRD complex (49%), with CoREST (28%) and SIN3 (15%) being the next most abundant. Collectively, these 3 complexes make up the majority of HDAC1 complexes in the cell at 92%. MIER (4%), MiDAC (3%) and RERE1 (1%) are minor species in ESCs, albeit, in the case of MiDAC, still essential for development ^12^. Reassuringly, the sum-total iBAQ value of HDAC1-binding proteins is 91% of the total HDAC1-Flag recovered in the pulldown, consistent with the 1:1 interaction between HDAC1 and its binding partners observed in multiple structural determinations ^12,15,16^. The estimated relative stoichiometry of individual complex components allowed a number of compelling insights. For example, the SIN3A complex and its core components (SIN3A, SUDS3 and SAP30) appear to be at least 10-fold more abundant in ESCs than the SIN3B complex (SIN3B, EMSY, ARID4B). The NuRD complex incorporates either methyl-CpG-binding domain protein 2 (MBD2) or MBD3 in a mutually exclusive manner ^24^. Here we show that MBD3 is clearly the major constituent in ESCs with a 10:1 MBD3/MBD2 ratio. Stoichiometric components of NuRD (based on MTA1-3 and MBD3 levels), include CHD4, GATA2A and GATA2B; while ZNF219, PARP1 and the helicase, CHD3 are only minor constituents. The core of CoREST consists of LSD1, RCOR1 or RCOR2 and HDAC1, with smaller amounts of ZMYM2. Notable CoREST-associated proteins, CTBP1 and CTBP2, appear to be sub-stoichiometric members of the complex. MIDEAS and TRERF1 are both ELM2-SANT-containing proteins that are likely interchangeable HDAC1-binders within MiDAC, with the former being the dominant component in ESCs. Both proteins interact with DNTTIP1, which was present in our ESC extracts, but did not reach significance following coIP (supplementary Table S1). RBBP4 and RBBP7 are well-known histone-binding subunits of multiple chromatin-modifying machines, including SIN3A, NuRD and PRC2 ^25–29^. We observed abundant levels of both proteins associated with HDAC1 and the RBBP4/RBBP7 ratio was 10:1 in our experiments (Fig 1D). Intriguingly, we measured almost twice as much RBBP4 as HDAC1, consistent with a 2:1:1 ratio of RBBP4/MTA1/HDAC1 in NuRD ^24^. Numerous HDAC1-associated transcription factors were also identified, most notably, SALL4, which is required for ESC pluripotency and is known to bind NuRD ^30–32^. Most of the other TFs identified were purified at levels far below those of the core complex components, which reflects both their relative expression levels and the transient nature of their association. We also observed a number of chromatin-modifying enzymes, including TET1 and OGT, known to bind to the SIN3A complex ^33,34^ and both EHMT1 (KMT1D/GLP) and EHMT2 (KMT1C/G9A), euchromatic histone H3 Lys9 methyltransferases, with roles in gene silencing.

**Figure 1.**
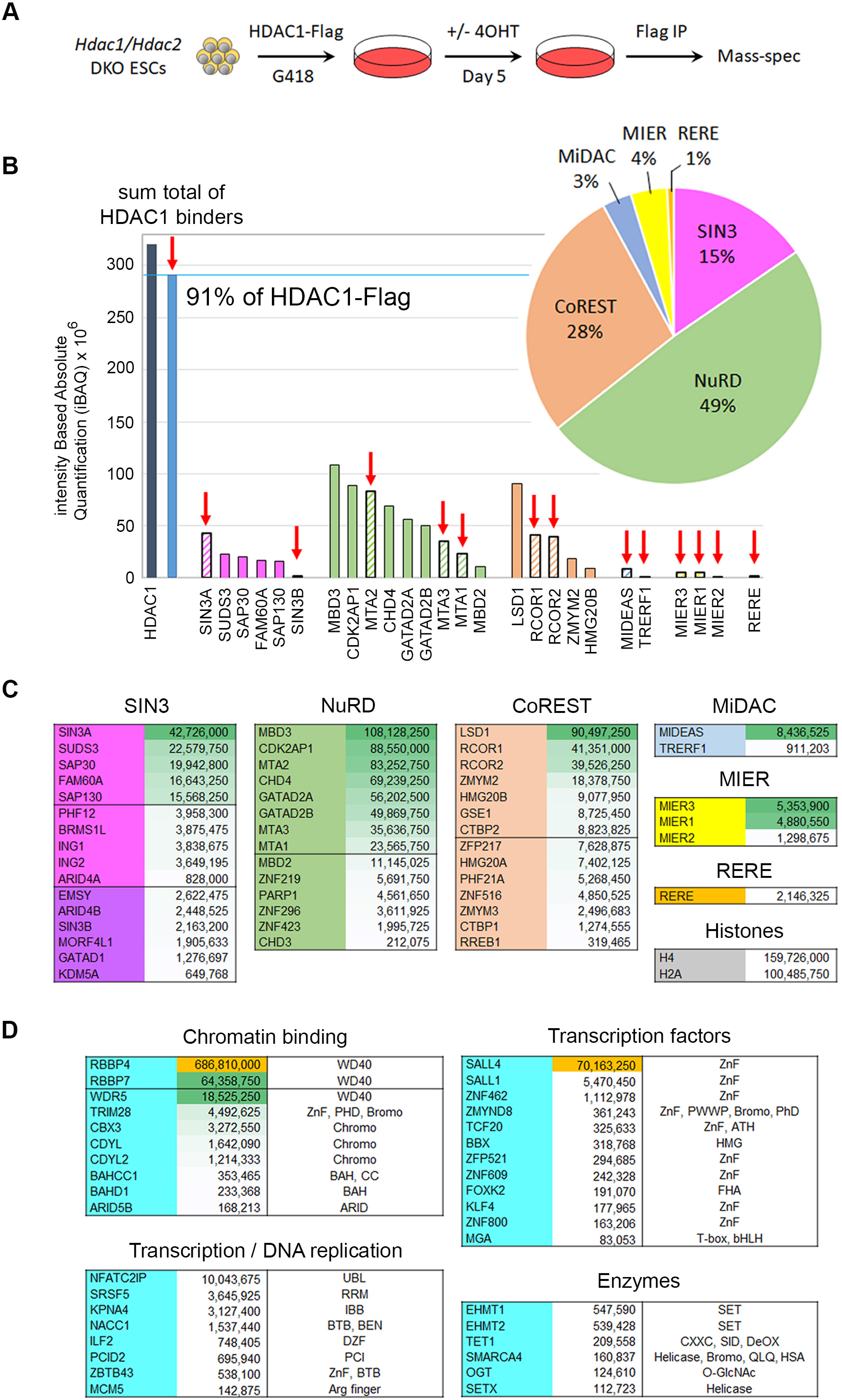
Relative abundance of HDAC1-binding proteins and associated complex components. (A) schematic showing experimental procedure to produce HDAC1-Flag co-immunoprecipitation mass spectrometry in *Hdac1/2* double knockout (DKO) cells. (B) Graph showing intensity-based absolute quantification (iBAQ) values for core components of the six unique HDAC1/2 complexes as indicated. The red arrow indicates proteins that directly bind to HDAC1. The pie chart indicates the relative proportion of the six HDAC1/2 complexes determined by the iBAQ values for direct HDAC1-binding components from each complex. (C) Table showing the iBAQ values for components of the indicated HDAC1/2 complexes ranked in order of abundance. (D) Table showing iBAQ values for significantly enriched HDAC1-associated proteins grouped into the indicated functional groups. Domains contained by these proteins are also indicated.

### ELM2/SANT and SIN3 proteins utilise distinct interacting surfaces on HDAC1

13 of 15 HDAC1 binding proteins utilise adjacent ELM2-SANT domains to directly interact with HDAC1 ^4,5^. The schematic diagram in Fig 2A (left) shows the effective arrangement of these two domains across the surface of HDAC1, based on high-resolution structures of MTA1:HDAC1 and MIDEAS:HDAC1 (Fig 2A, right)^12,15^. The SANT domain consists of 3 α-helices, which together with a positively-charged surface on HDAC1, sandwiches a molecule of inositol phosphate (InsP) necessary for full enzymatic activity ^35^. The ELM2 domain has a bipartite structure, with an unstructured N-terminal region that lies across a highly conserved groove on the underside of HDAC1, and a C-terminal region containing a 3 α-helical bundle that sits adjacent to the SANT domain. These extensive interactions can be subdivided into 3 discrete regions (Fig 2A, see left panel): SANT (1), ELM2-C (2) and ELM2-N (3). To identify residues on the surface of HDAC1 required for interaction with the 6 unique multiprotein complexes, we used the MTA1/HDAC1 structure to design mutations in regions 1, 2 and 3 to perturb binding. In region 1, we focussed on a pair of Tyr residues, Y333/Y336 (Fig 2B), that mediate interactions with phosphates of the InsP6 and residues in α3 of the SANT domain. We mutated Y48 in region 2 which forms interactions with α2 of the ELM2-C domain. Region 3 forms an extended surface over which the unstructured ELM2-N domain stretches, forming a variety of interactions (Fig 2B). We therefore chose 3 charged (E63R, K126E and K144E), 1 hydrophobic (L161E) and 1 aromatic/non-polar (Y166E) mutations that span both the breadth and diversity of this region. Initially, we tested the effects of these mutations on HDAC1 when expressed and purified from HEK293T cells (supplementary Fig S1). HDAC1 expression levels were unaffected by the mutations, however, we observed significant changes in their relative deacetylase activity. HDAC1 Y333A/Y336A, K126E and L161E/Y166R all showed activity below 10% of wild-type. Since none of these residues are adjacent to the active site, we infer residues on the surface of HDAC1 interacting with both SANT (region 1) and ELM2-N (region 3) contribute indirectly to enzymatic activity. In contrast, HDAC1 Y48E and E63R showed activity close to that of the wild-type enzyme. To monitor the interaction of wild-type and mutant HDAC1 with SIN3A we expressed both proteins transiently and performed a Flag-coIP, followed by western blotting. Intriguingly, we found a robust interaction between exogenous HDAC1-Flag and Myc-SIN3A proteins, but not endogenous SIN3A (supplementary Fig S1, compare red and green bands in the input vs coIP), suggesting that there is little interchange between corepressor and HDAC1 once the complex is assembled in cells. Mutation of Y333/Y336 to Ala or Asp reduced binding to all complexes, demonstrating the necessity of residues within region 1. In contrast, Y48E in region 2, bound to SIN3A while abolishing any interaction with RCOR1 (supplementary Fig S1 – middle and lower panels). Thus, demonstrating for the first time, distinct surface requirements between HDAC1-binding proteins.

**Figure 2.**
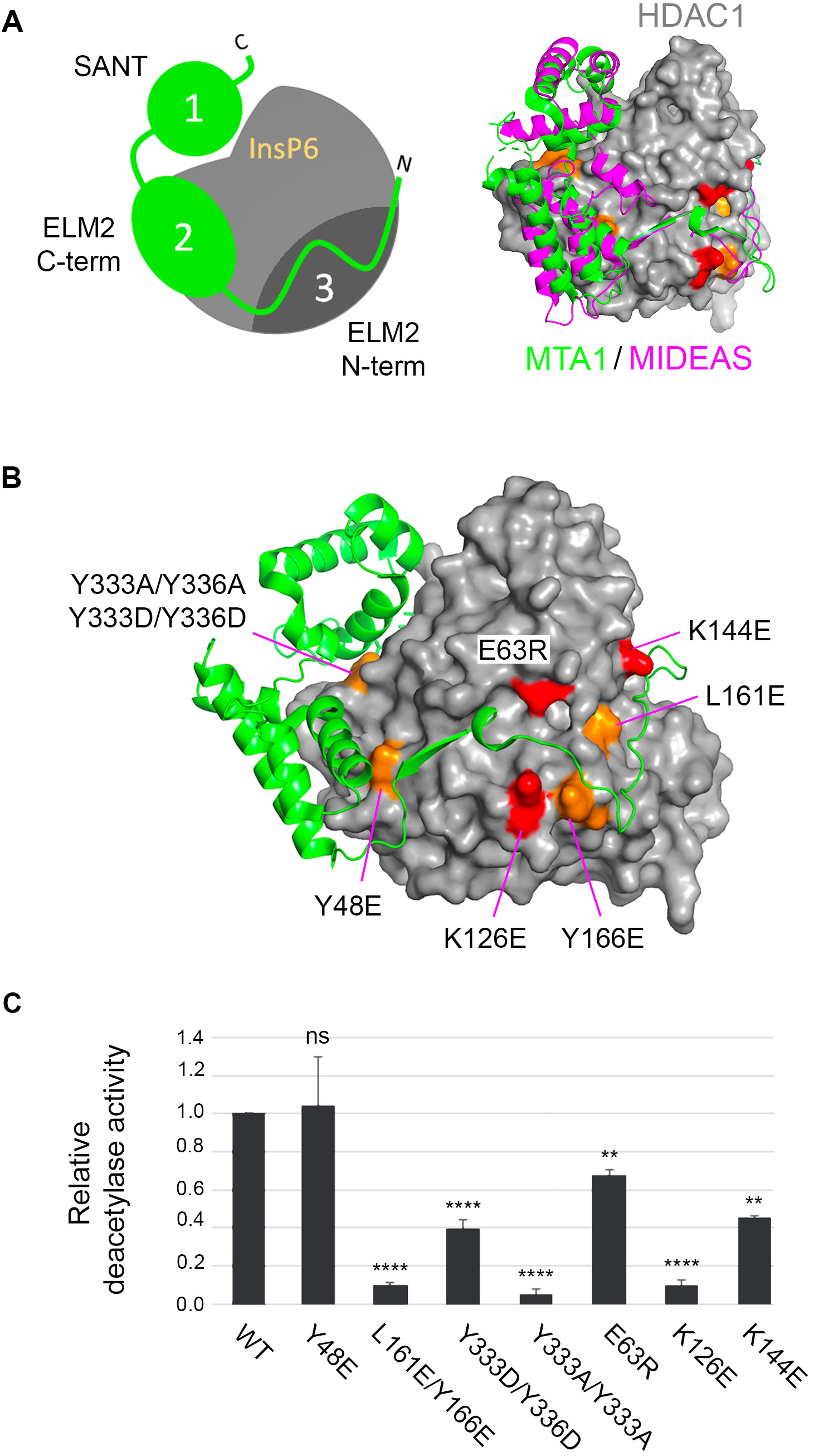
Defining critical interactions on the surface of HDAC1. (A) left panel, a schematic diagram outlining 3 separate regions used by ELM-SANT domain-containing HDAC1-binding proteins. The right panel, shows an overlay of MTA1/HDAC1 and MIDEAS/HDAC1 binary complexes, showing a conserved mechanism of HDAC1 binding. (B) HDAC1/MTA1 binary showing the position of individual residues mutated in the study. Orange, aromatic/non-charged residues; red, charged residues. (C) The deacetylase assay was performed using enzymes co-immunoprecipitated from HEK293T cell transfected with either wild-type HDAC1 or the indicated mutants. A one-way ANOVA was performed compared to WT values. NS, not significant, p >0.05, ** ≤ 0.01, **** ≤ 0.0001.

To examine the HDAC1 mutants in a more physiological environment, we used the piggyBAC system to stably introduce them into *Hdac1/2* DKO ESCs, so that following removal of the endogenous proteins (5 days post 4OHT), we could assess their capacity to rescue cell viability and interact with all 6 complexes (Fig 3A). We examined the survival of DKO cells expressing either WT or mutant HDAC1 using crystal violet staining (Fig 3B) and CellTiter-Glo (Fig 3C). Both assays revealed that expression of wild-type HDAC1 and E63R recovered cell viability to control levels, while Y333A/Y336A, K126E and L161E/Y166E (which all displayed severely reduced HDAC activity) were incapable of rescuing the cell death phenotype. Intriguingly, cells expressing HDAC1-Y48E, which only bound SIN3A in HEK293T experiments, showed approximately 50% cell viability in DKO ESCs, suggesting that retention of SIN3A binding alone is sufficient to retain some level of cell viability. Since HDAC1-Y48E and E63R mutants produced viable cells, we took these forward and performed large-scale coIPs for western blotting (Fig 3D) and mass spectrometry (Fig 3E) using DKO cells lacking a Flag epitope as a control (n = 4). WT HDAC1-Flag pulled down endogenous SIN3A, LSD1, MTA1 and DNTTP1 from ESCs, thus showing incorporation into the SIN3, CoREST, NuRD and MiDAC complexes respectively (Fig 3D). In contrast, the Y48E mutation bound only to SIN3A, while disrupting binding to all ELM2-SANT-containing partners (LSD1, MTA1 and DNTTIP1). HDAC1-E63R bound SIN3A and LSD1, but not MTA1 or DNTTIP1, thus demonstrating the ability to differentiate between individual ELM2-SANT-containing complexes.

**Figure 3.**
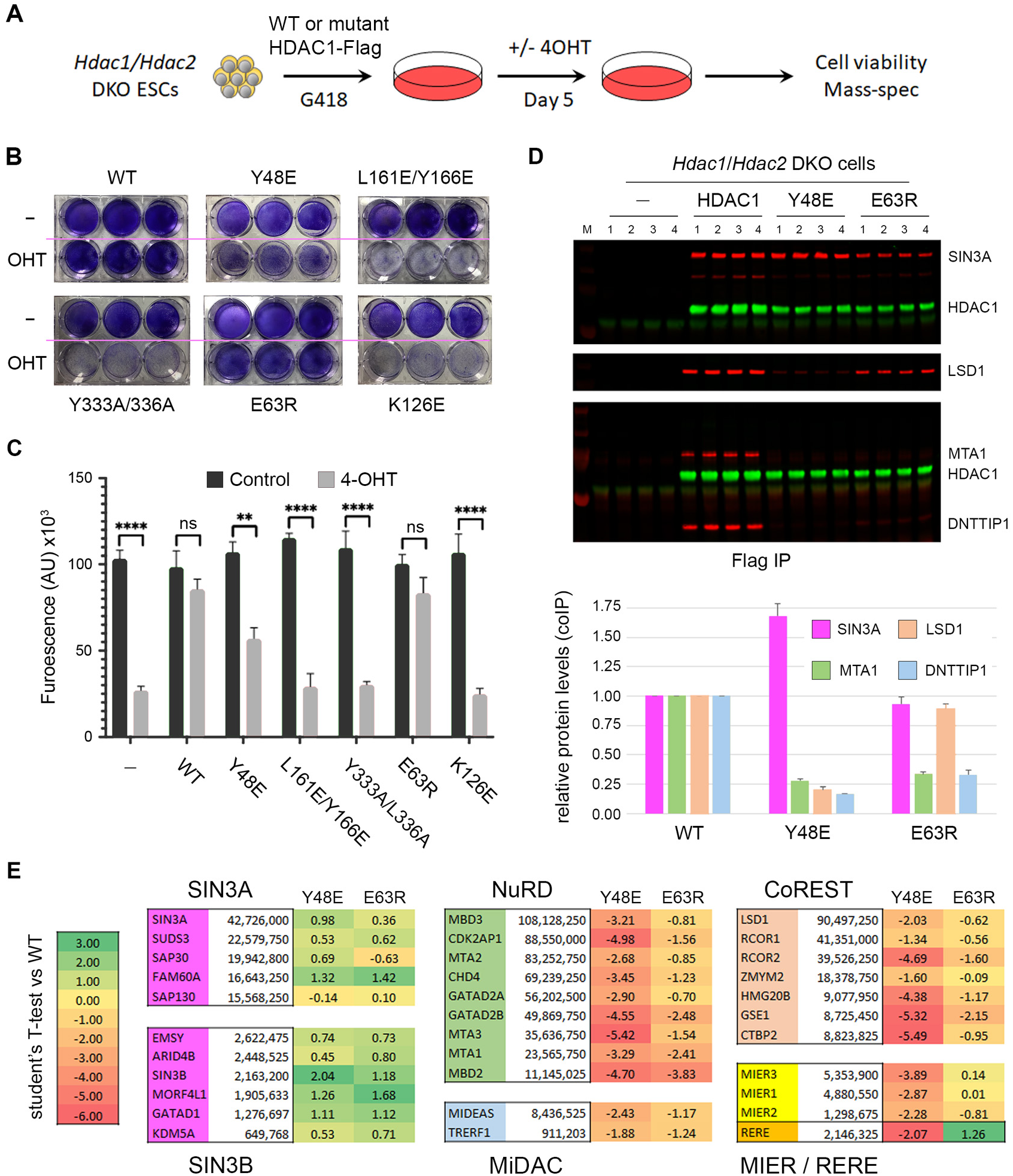
Individual point mutations discriminate between HDAC1-binding proteins. (A) schematic showing experimental procedure to reintroduce HDAC1-Flag, or the indicated mutants to *Hdac1/2* double knockout (DKO) cells for cell viability and coIP/mass spectrometry experiments. The ability of wild-type (WT) and mutant HDAC1 to rescue the viability of HDAC1/2 DKO cells using (B) crystal violet staining of cells at 5 days post 4OHT treatment, and (C) a CellTiter-Glo (CTG) assay. (D) top panel, western blot for the indicated proteins following Flag-IP from DKO cells expressing either WT or Y48 and E63R mutations. Lower panel, quantification of the indicated proteins from western blot data normalised to the amount of HDAC1 in the coIP. Mean values for biological replicates (n=4) are plotted with error bars indicating standard deviation from the mean. (E) The relative interaction of individual HDAC1/2 complex components was determined for Y48E and E64R mutants using mass spectrometry from n=4 biological replicates. Values indicate Log_2_ student’s T-test of the indicated mutant versus binding to wild-type HDAC1. Positive values show increased binding and negative values a relative reduction in binding compared to control.

To extend this analysis to all complexes and their components, we used material from the Flag-coIP to identify proteins bound to WT HDAC1, Y48E and E63R via mass spectrometry (Fig 3E). We performed statistical analysis (student’s T-test permutation-based FDR 0.05, WT vs mutant) to evaluate changes in the binding of core components from all six HDAC1/2 containing complexes to either Y48E or E63R. HDAC1-Y48E showed increased binding to all but one component of the SIN3A and SIN3B complexes (10 out of 11), but a strong reduction to all components of ELM-SANT dependent complexes (22 out of 22). Thus, mutation of a single residue on the surface of HDAC1 in region 2 can abolish binding to 5 of 6 HDAC1/2 complexes. HDAC1-E63R also showed increased binding to SIN3A/B complexes and could further distinguish between ELM-SANT complexes. MIER1 and MIER3 showed no change in binding between WT and E63R, while RERE binding was slightly increased. In contrast, we observed reduced binding to all components of the NuRD and MiDAC complexes consistent with the absence of DNTTIP1 in western blots (Fig 3D). We have thus identified two separate regions on the surface of HDAC1 capable of distinguishing incorporation into specific complexes. Y48E presents a black-and-white view of binding with disruption of all ELM2/SANT partners, retaining only SIN3A/B. HDAC1-E63R on the other hand, retains SIN3A, MIER1, RERE and residual binding to CoREST, while abolishing the association of NuRD and MiDAC.

### Retaining only a subset of HDAC1/2 complexes leads to histone hyperacetylation and loss of transcriptional regulation

HDAC1-Y48E and E63R retain HDAC activity (Fig 2C) and the ability to rescue DKO ESC viability (Fig 3B and 3C), but only bind to a limited number of HDAC1-interacting partners (Fig 3D and 3E). Since only a subset of HDAC1/2 complexes remain active in these cells we hypothesised that this would cause changes in both histone acetylation and gene regulation. To examine this, we isolated histone proteins from control and HDAC1 mutant cells and then measured changes in acetylation levels by mass spectrometry (supplementary Table S3). HDAC1-Y48E, which only binds SIN3A/B, caused significant increases in acetylation at multiple lysine residues within H2B, H3 and H4 compared to WT controls (Fig 4A). Specific sites in H2B (K12, K15 and K20) and H3 (K14, K18 and K23) showed the greatest increase in acetylation. Interestingly, these same sites are also sensitive to loss of p300/CBP activity ^2^, suggesting an interplay of HDAC1/2 complexes with these critical HAT enzymes. HDAC1-E63R, which retains binding to SIN3A/B, CoREST, MIER and RERE, and is therefore the weaker of the two mutants tested, showed more subtle changes in histone acetylation. Although we could detect increases in acetylation of H2B and H3 these did not reach statistical significance. Overall, this is consistent with E63R being incorporated into more HDAC1/2 complexes compared to Y48E.

**Figure 4.**
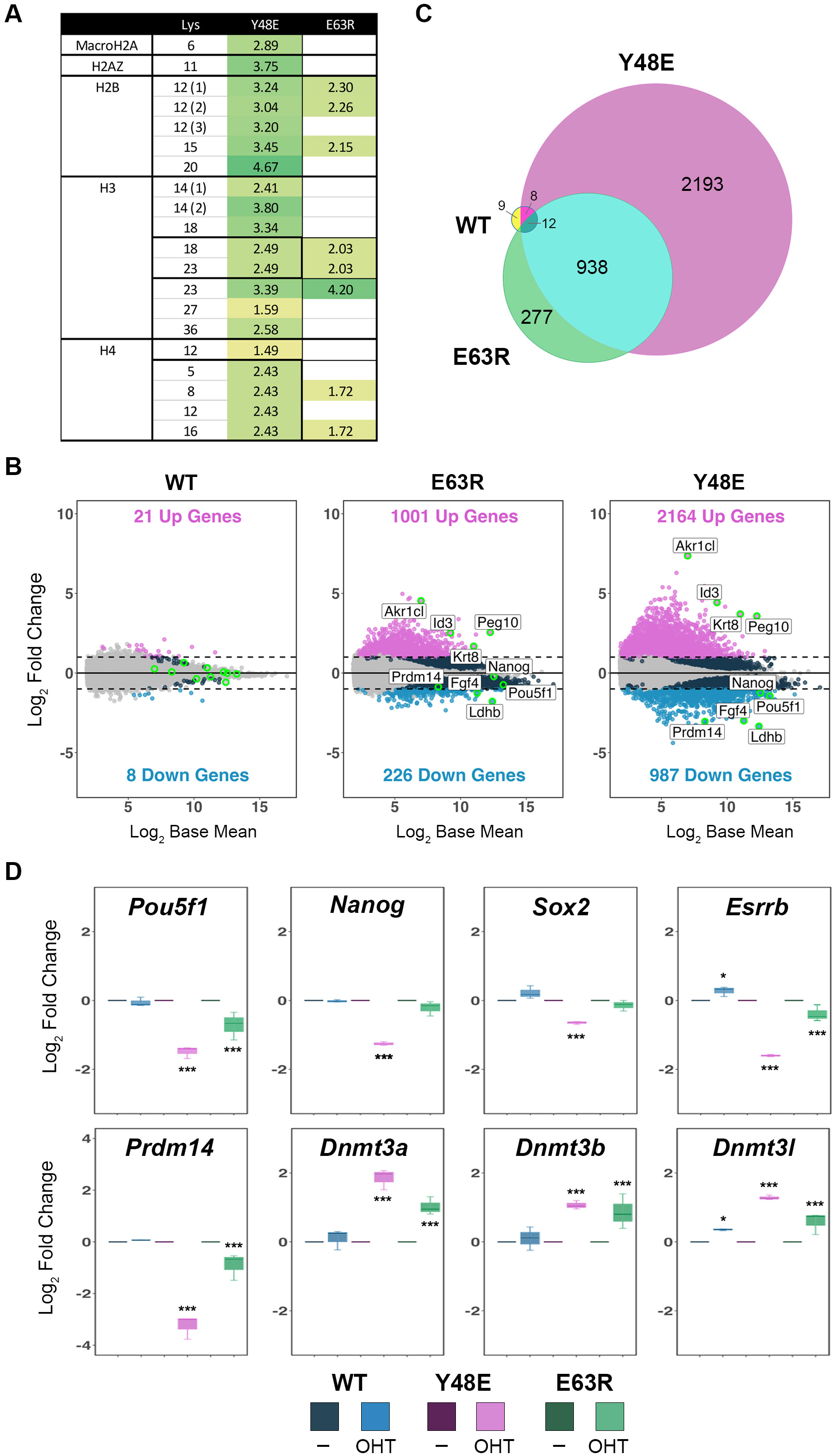
HDAC1 mutants that disrupt complex incorporation cause histone hyperacetylation and transcriptional dysregulation. (**A**) change in acetylation relative to wild-type is shown for the indicated Lys residues, determined by mass spectrometry from n=4 biological replicates. Brackets indicate measurements from unique peptides. Bold lines show multiple acetylation sites from the same peptide. Values are Log_2_ student’s T-test values for each mutant versus WT controls. (**B**) MA plots indicate differentially expressed genes (DEGs) for *Hdac1/2* DKO cells rescued with WT or E63R and Y48E mutants from n=3 biological replicates. Cut-offs applied were >2-fold change in expression, p-adjusted value <0.01. (**C**) Venn diagram showing the number and overlap of DEGs from WT, E63R and Y48E expressing cells. (**D**) Boxplots displaying relative fold-change for the indicated genes. The adjusted p-values were taken from DESeq2 using a Wald test and the Benjamini and Hochberg method to correct for multiple hypothesis testing. * p<0.01, ** p<0.001, *** p<0.0001.

We next examined the transcriptome of *Hdac1/2* DKO ESCs rescued with either HDAC1-WT or Y48E and E63R mutants (Fig 4B). The addition of HDAC1-WT to DKO cells completely rescues the gene expression phenotype, with only minor changes in transcription (21 genes up and 8 down). In contrast, HDAC1-E63R produced significant numbers of differentially expressed genes (DEGs, 1001 up and 226 down). Consistent with its stronger effect on binding, the Y48E mutation produced a greater number of DEGs (2164 up and 987 down) (see supplementary Table S2 for a complete list of DEGs). There is a clear correlation between the number of functional HDAC1/2 complexes and the number of dysregulated genes. We observed a significant overlap of genes between the two mutants, with 76% of DEGs in the E63R mutant coinciding with Y48E (Fig 4C, 938 of 1227). Gene ontology analysis of overlapping DEGs from Y48E and E63R mutants identified significant changes in multiple pathways, including, placental development, vasculogenesis and keratinization. A number of keratin genes showed significant de-repression, including *Krt5* (31-fold), *Krt14* (26-fold), *Krtdap* (9-fold) and *Krt8* (3-fold). Although we observed a good overlap between the two mutants, the majority of DEGs in Y48E cells were unique to that mutation (70%, 2201 of 3151). Among the unique pathways perturbed by HDAC1-Y48E was the cellular response to leukaemia inhibitory factor (LIF), a critical signalling pathway for the maintenance of pluripotency in mouse ESCs. Upon closer examination, we observed down-regulation of multiple pluripotency-associated transcription factors, including, *Pou5f1*, *Nanog* and *Esrrb* (Fig 4D). ESCs have relatively low levels of DNA methylation, in part due to the repression of DNA methyltransferase (DNMT) enzymes by PRDM14 ^36,37^. The HDAC1-Y48E cells showed a decrease in PRDM14, consistent with a loss of pluripotency, and a corresponding increase in *Dnmt3a*, *Dnmt3b* and *Dnmt3l* levels. These data demonstrate the requirement for a complete array of HDAC1/2 complexes for normal transcriptional regulation and homeostasis in ESCs.

### Comparative analysis of HDAC1 binding partners reveals distinct interactions with a conserved interaction surface

To understand better the binding properties of Y48E and E63R mutants and the molecular determinants of HDAC1 interaction more generally, we performed a comparative analysis using binding partners from all 6 complexes. These include a recent cryoEM structure of SIN3B/HDAC2 ^17^, NuRD ^12^ and MiDAC ^24^ complexes, as well as AlphaFold3 ^38^ models of MIER1 and RERE bound to HDAC1. What is immediately obvious, is that the mechanism of HDAC recruitment into the SIN3B complex is profoundly different to those complexes employing ELM2/SANT domains (Fig 5A). SIN3B binding is largely confined to the surface of HDAC1 in proximity to the active site, opposite to Y48 and E63, explaining why these mutations have little effect on SIN3 binding (compare front vs back of HDAC1, supplementary Fig S3). In contrast, the ELM2-SANT domain is in direct contact with Y48. A conserved acidic residue is incompatible with the Y48E mutation. There are however binding surfaces conserved between SIN3 and ELM2-SANT binders, including Y333/Y336 (Fig 5A), such that double mutation to Ala or Asp prevents both types of interaction (supplementary Fig S1). The HDAC surface around E63 provides an area of distinctiveness between ELM2-SANT binding partners. We found that MIER1 and RERE were unaffected or showed a slight increase in binding to the E63R mutation, respectively. Interaction with RCOR1 and RCOR2 from the CoREST complex was reduced, but western blotting clearly shows a residual interaction (Fig 3D). All components NuRD and MiDAC were reduced (Fig 3E). Examining the molecular details of these interactions, we identified conserved Arg (R189 in MTA1 and R130 in RCOR1) and Lys (K748 in MIDEAS) residues positioned directly towards E63, explaining why these 3 complexes are most affected by this mutation (Fig 5B, 5C). The unstructured ELM2-N loop of RERE appears to circumvent E63 altogether, which explains the increased binding to the E63R mutant (Fig 3E), particularly in the absence of competing HDAC1-binders. MIER1 associates with the HDAC1-E63 pocket using a non-polar interaction with Y206 and is thus unaffected by the switch in charge (Fig 5C). By exploiting the unexpected sensitivity of HDAC1-binding proteins to specific surface residues we are now able to differentiate between the recruitment of these crucial histone-modifying machines.

**Figure 5.**
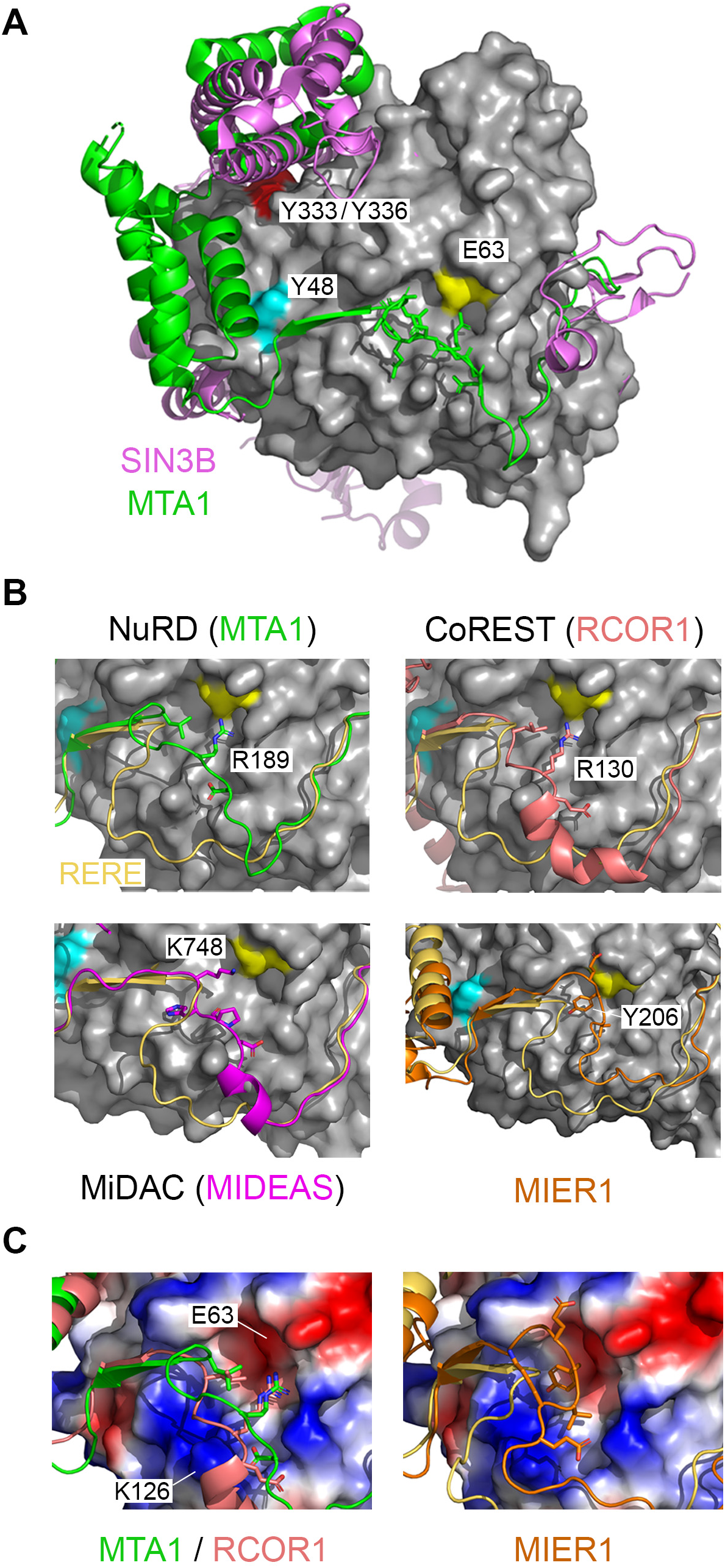
HDAC1 binding proteins have distinct modes of interaction. (**A**) binary structures of MTA/HDAC1 (5ICN) and SIN3B/HDAC2 (8BPA) are superimposed to demonstrate different binding modalities. Residues required for interaction with binding partners are labelled. (**B**) a binding pocket centred on HDAC1-E63 (yellow) displays conserved and distinct interactions with ELM2/SANT-containing partners as indicated. An AlphaFold3 model of HDAC1/RERE (gold) is used as a reference point for other ELM2/SANT interactors. (**C**) shows the electrostatic surface around HDAC1-E63 utilised by MTA1 and RCOR1, in contrast to MIER1.

## Discussion

The majority of histone-modifying enzymes operate in multiprotein complexes with, PRC2 ^39^, SAGA ^40^ and SET1 ^41^ providing well-studied paradigms. There is though, perhaps no greater variety of complexes housing a single enzymatic activity than the HDAC1/2 deacetylase complexes. HDAC1/2 are incorporated into six unique complexes (SIN3A/B, NuRD, CoREST, MiDAC, MIER and RERE) as defined by multiple previous coIP/mass spectrometry studies ^42–46^. Why the cell requires so many different vehicles for HDAC1/2 activity remains to be fully understood. While there have been extensive mapping studies, here we sought to address the possibility of a hierarchy among individual HDAC1/2 complexes. CoIP of HDAC1-Flag from HDAC1/2 DKO cells showed that there are 3 dominant HDAC1/2 complexes, NuRD (49%), CoREST (28%) and SIN3A (15%), constituting 92% of the total HDAC1 complexes in ESCs. The remaining complexes, MiDAC (3%), MIER (4%) and RERE (1%) make up a relatively minor cohort. Abundance does not entirely define importance of course, deletion of either of the MiDAC components, *Dnttip1* or *Mideas*, caused embryonic lethality in the mouse at e16.5 days ^12^. However, KO studies involving components of the NuRD, SIN3 and CoREST all produced early embryonic phenotypes prior to gastrulation, perhaps hinting at a greater importance at earlier time points ^6^. Our experiments also provide insights into the relative composition of the different HDAC1/2 complexes. SIN3A appears to be ten-fold more abundant than SIN3B for instance (Fig 1B). Similarly, the ratio of MBD3 vs MBD2 is 10:1, which may have implications for their distinct activities in the NuRD complex. These data also give an indication of each of the *core* complex components and those present in sub-stochiometric quantities, CHD4 vs. CHD3 (a 325:1 ratio) in the NuRD complex, or CTBP2 vs. CTBP1 (an 8:1 ratio) in the CoREST complex, being two of many examples (Fig 1C). To gain an understanding of the consistency of these ratios between cell types, we compared our data to the HDAC1 coIP reported by Vcelkova and colleagues ^45^ (shown in supplementary Fig S4). Consistent with our study, NuRD was also the most abundant complex by far in mouse embryo fibroblasts (MEFs) and HAP1 cells. However, the ratio of MBD3/MBD2 in MEFs is approximately 1:1, compared with 10:1 in mouse ESCs. The MiDAC complex also appears to be similar in abundance to SIN3A and CoREST in MEFs (supplementary Fig S4). These data suggest that the relative level of HDAC1/2 complexes and their components may differ somewhat depending on cell type and require further investigation.

While there have been numerous HDAC1 mutations made previously, the bulk of these have focussed on the catalytic site of the enzyme ^47–49^. We present here, the first systematic study of HDAC1 surface residues required to bind 15 separate interacting proteins from each of the 6 complexes. 13 of these proteins utilise a combination of ELM2-SANT domains to bind to HDAC1, superficially at least, a cut-and-paste mechanism utilised by nature for interaction with HDAC1. Using a comparative analysis of HDAC1/MTA1 and HDAC1/MIDEAS structures ^12,15^ we identified 3 separate regions utilized by these binary complexes (Fig 2A). We hypothesized that loss of binding in any one of these regions would be *insufficient* to disrupt HDAC1 interaction since there would still be binding to the remaining two. To our surprise, we found that single (Y48E, K126E) and double (L161E/Y166E, Y333A/Y336A) mutations within each of these regions were capable of disrupting binding to all ELM2/SANT domain-containing proteins (Fig 3 and supplementary Fig S1). Interestingly, Y333A/Y336A (in region 1), whose aromatic rings stack to support interactions with the SANT domain, abolished binding to all HDAC1-binders, including SIN3A/B. Y48 (in region 2) occupies a position on the surface that interacts with the dimerization domain of MTA1 (Fig 2B ^15^). Mutation to glutamic acid (Y48E) caused loss of binding to all ELM2-SANT domain binders, but retention of binding SIN3A/B. When we introduced HDAC1-Y48E into ESCs, we monitored *increased* binding to SIN3A/B complex components to the mutant (Fig 3E), presumably because no other factors were competing for HDAC1 binding. Remarkably, HDAC1-Y48E was able to partially rescue the viability of HDAC1/2 DKO cells (∼50% cell viability), suggesting that retention of the SIN3A/B complexes is sufficient for ESC viability (Fig 3C). We identified a second more subtle mutation in region 3 (E63R), that has near wild-type deacetylase activity and ability to rescue DKO cells, that discriminates between the different ELM-SANT complexes (Fig 2B and Fig 3D).

There is an assumption that most, if not all, HDAC1/2 complexes play a role in gene regulation. This has been demonstrated comprehensively for the 3 dominant complexes, NuRD, CoREST and SIN3A, which are all recruited to DNA by a variety of transcription factors ^6,50^. The jury is still out in this regard for MiDAC and MIER, which have putative roles in mitosis ^12,43^ and as a histone chaperone ^51^ respectively. Cells expressing HDAC1-Y48E retain only functional SIN3A and SIN3B complexes, which is sufficient to retain viability in ESCs, but results in global histone hyperacetylation and the dysregulation of over 3,000 genes (Fig 4A and Fig 4B). The scale of the effect provides a potent example of the necessity for a full repertoire of ELM2-SANT domain containing complexes. HDAC1-E63R is the first mutation to discriminate between the binding of ELM2-SANT-dependent HDAC1-binding proteins (Fig 3D). E63R bound MIER1-3 and RERE at the same level as wild-type while perturbing binding to MTA1-3, RCOR1/2 and MIDEAS (Fig 3E). These data demonstrate for the first time that despite a common HDAC1 binding modality, the precise molecular interactions on the surface of the enzyme are distinct. In theory, at least, this might suggest that we could exploit differential binding to disrupt protein-protein interactions within each complex and thus generate a complex-specific HDAC inhibitor. In this study, we have defined critical residues on the surface of HDAC1 necessary for incorporation into a diverse group of multiprotein complexes and that against expectation, single point mutations are able to distinguish between the binding of even closely related ELM2/SANT-containing proteins. Consequently, different patterns of binding led to differential gene expression highlighting the distinct function of HDAC1/2 complexes in cells and development.

## Acknowledgments

Thank you to members of the Cowley and Schwabe groups for critical comments and feedback on the data throughout the project. We thank Dr. Nicolas Sylvius and the NUCLEUS genomics facility for help with RNA extractions and quality checking of RNA/DNA. Plasmid constructs were generated by the PROTEX facility at the University of Leicester. This research used the ALICE High Performance Computing facility at the University of Leicester

---

**Fig S1.**
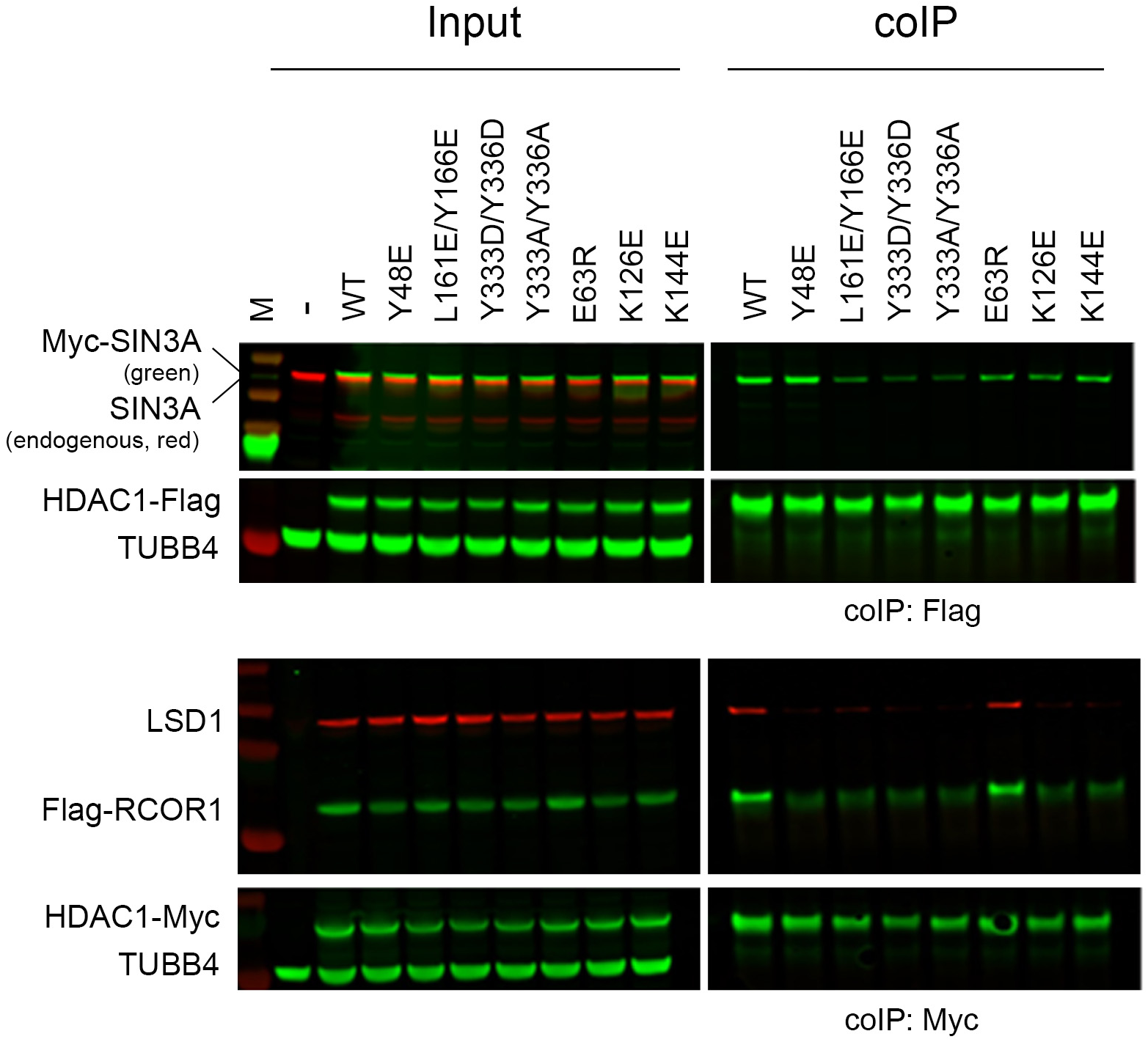
HDAC1 surface mutations show differential binding to corepressor partners. Wildtype or mutant HDAC1 with a Flag-tag or Myc-tag (as indicated) were immunoprecpitated from HEK293T cells. Western blot shows input or binding (colP) of the indicated proteins. TUBB4 was used as a loading control.

**Fig S2.**
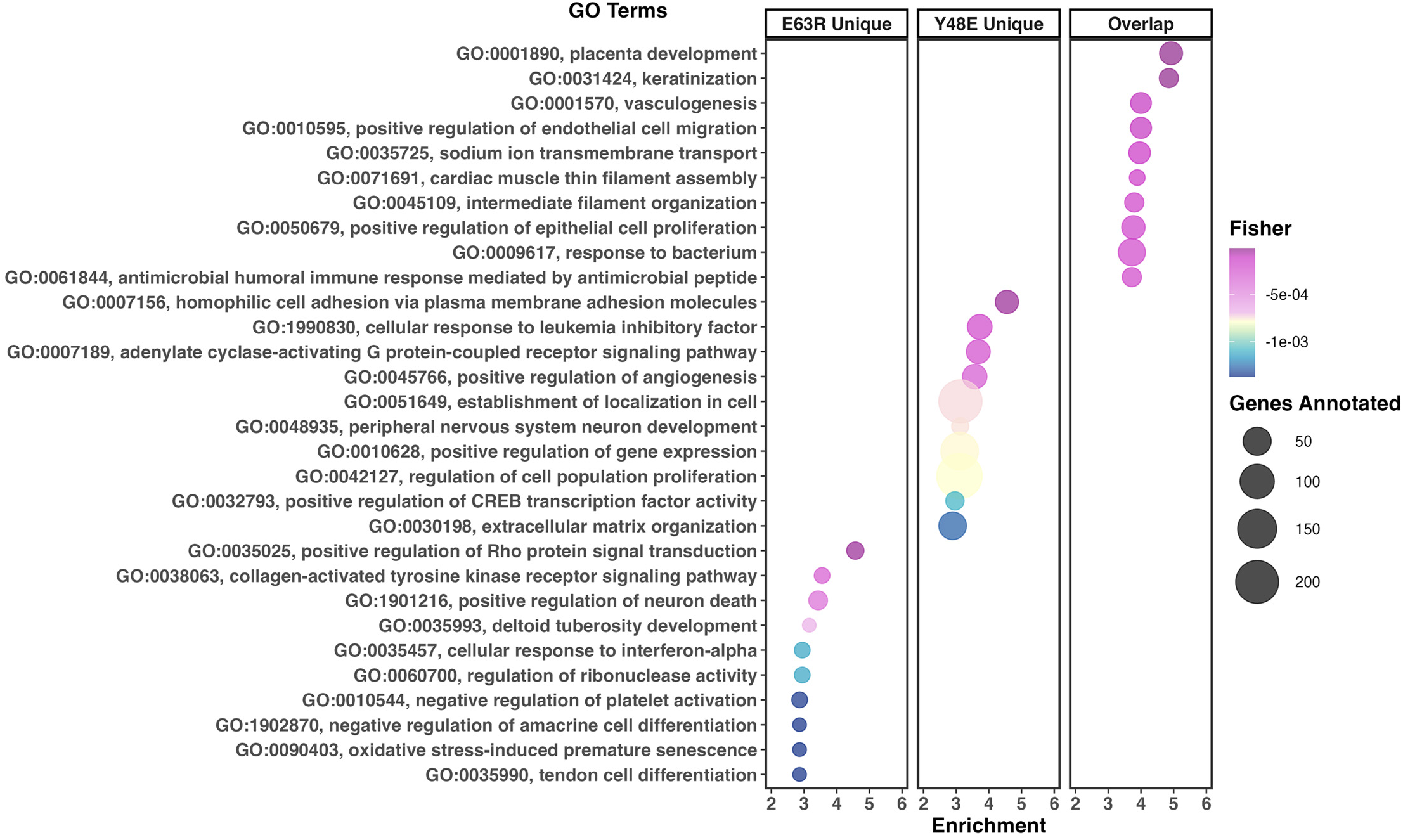
Gene ontology (GO) analysis of differentially expressed genes (DEGs). The 10 most enriched biological process GO terms associated with each individual mutant *(E63R unique* and *Y48E unique)* or shared *(overlap)* are shown (padj <0.01, log_2_ fold change> −1) following 24 hours of HDAC1-dTAG degradation.

**Fig S3.**
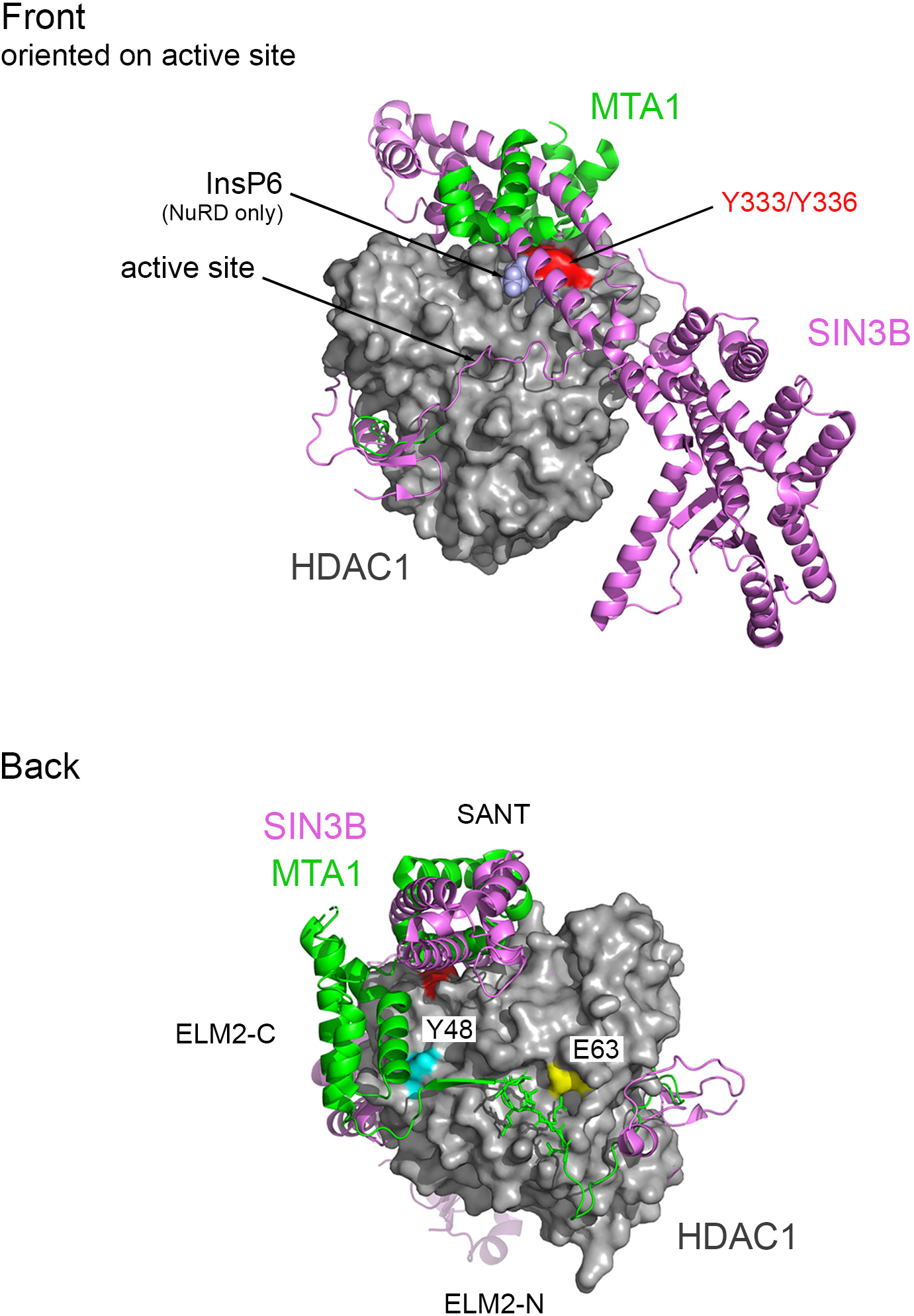
HDAC1 binding proteins have distinct modes of interaction. Binary structures of MTA/HDAC1 (5ICN) and SIN3B/HDAC2 (8BPA) are superimposed to demonstrate different binding modalities. Residues required for interaction with binding partners are labelled. The front of HDAC1 (based on the position of the active site, as indicated) is shown above, with the back below. lnsP6 is present only in the NuRD complex, but has been included as a reference point.

**Fig S4.**
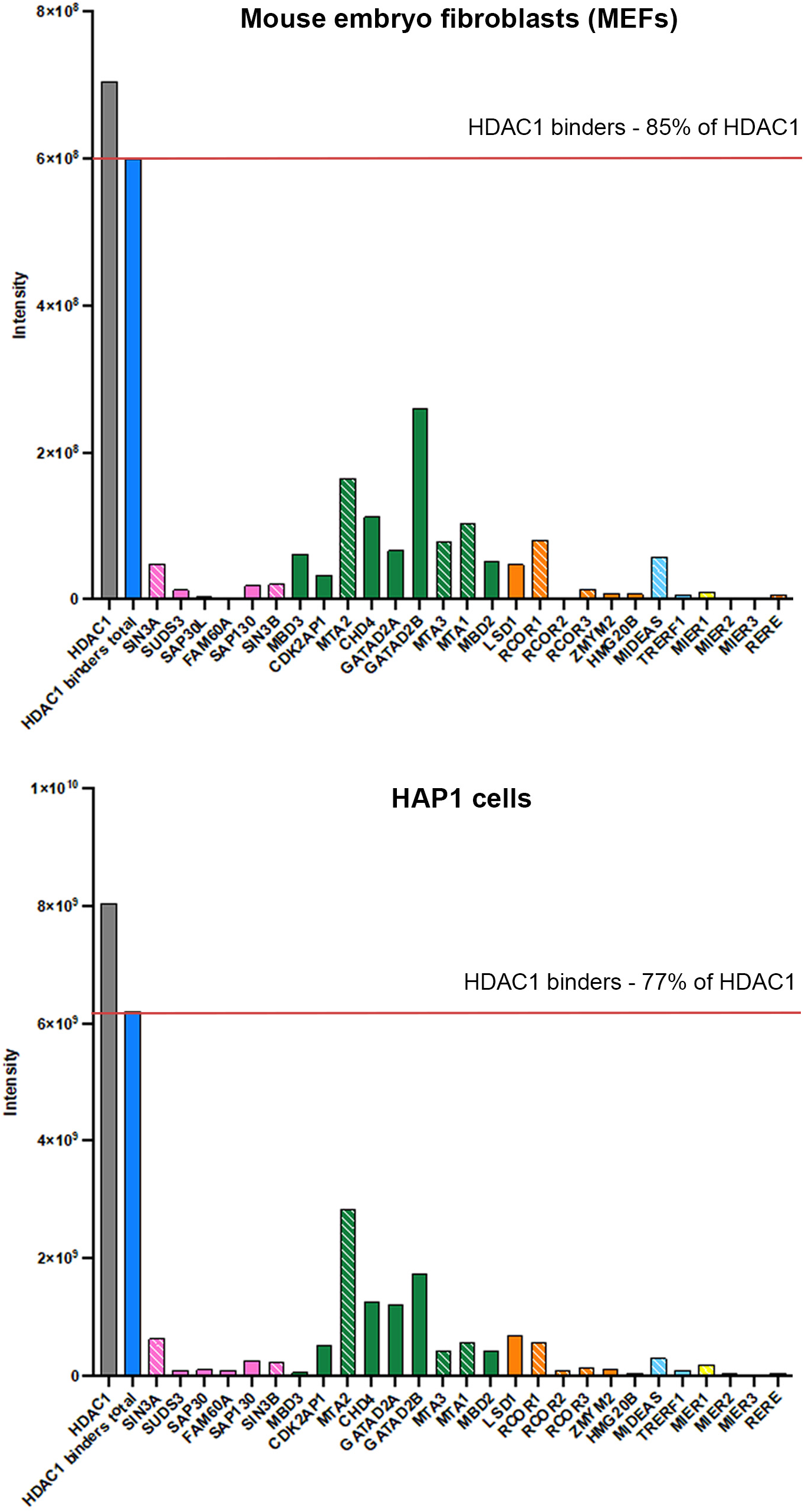
HDAC1 co-immunoprecipitation / mass-spectrometry data taken from Vcelkova et al., 2023 (PMID:37878419). Intensity values for individual components of HDAC1/2 containing complexes are shown from MEFs and HAP1 cells. Direct HDAC1 binding proteins are indicated by hashed lines.

## References

1. Barnes, C.E., English, D.M. & Cowley, S.M. Acetylation & Co: an expanding repertoire of histone acylations regulates chromatin and transcription. Essays Biochem 63, 97–107 (2019).

2. Weinert, B.T. et al. Time-Resolved Analysis Reveals Rapid Dynamics and Broad Scope of the CBP/p300 Acetylome. Cell 174, 231–244 e12 (2018).

3. Zheng, Y., Thomas, P.M. & Kelleher, N.L. Measurement of acetylation turnover at distinct lysines in human histones identifies long-lived acetylation sites. Nat Commun 4, 2203 (2013).

4. Millard, C.J., Watson, P.J., Fairall, L. & Schwabe, J.W. Targeting Class I Histone Deacetylases in a “Complex” Environment. Trends Pharmacol Sci 38, 363–377 (2017).

5. Asmamaw, M.D., He, A., Zhang, L.-R., Liu, H.-M. & Gao, Y. Histone deacetylase complexes: Structure, regulation and function. Biochimica et Biophysica Acta (BBA) - Reviews on Cancer 1879, 189150 (2024).

6. Kelly, R.D. & Cowley, S.M. The physiological roles of histone deacetylase (HDAC) 1 and 2: complex co-stars with multiple leading parts. Biochem Soc Trans 41, 741–9 (2013).

7. Cowley, S.M. et al. The mSin3A chromatin-modifying complex is essential for embryogenesis and T-cell development. Molecular and cellular biology (2005).

8. Dannenberg, J.-H. et al. mSin3A corepressor regulates diverse transcriptional networks governing normal and neoplastic growth and survival. Genes & development 19, 1581–1595 (2005).

9. David, G. et al. Specific requirement of the chromatin modifier mSin3B in cell cycle exit and cellular differentiation. Proc Natl Acad Sci U S A 105, 4168–72 (2008).

10. Jelinic, P., Pellegrino, J. & David, G. A novel mammalian complex containing Sin3B mitigates histone acetylation and RNA polymerase II progression within transcribed loci. Molecular and cellular biology 31, 54–62 (2011).

11. Hendrich, B., Guy, J., Ramsahoye, B., Wilson, V.A. & Bird, A. Closely related proteins MBD2 and MBD3 play distinctive but interacting roles in mouse development. Genes & development 15, 710–723 (2001).

12. Turnbull, R.E. et al. The MiDAC histone deacetylase complex is essential for embryonic development and has a unique multivalent structure. Nat Commun 11, 3252 (2020).

13. Wang, J. et al. The lysine demethylase LSD1 (KDM1) is required for maintenance of global DNA methylation. Nature genetics 41, 125–129 (2009).

14. Wang, J. et al. Opposing LSD1 complexes function in developmental gene activation and repression programmes. Nature 446, 882–887 (2007).

15. Millard, C.J. et al. Class I HDACs share a common mechanism of regulation by inositol phosphates. Mol Cell 51, 57–67 (2013).

16. Song, Y. et al. Mechanism of crosstalk between the LSD1 demethylase and HDAC1 deacetylase in the CoREST complex. Cell Reports 30, 2699–2711. e8 (2020).

17. Wan, M.S. et al. Mechanism of assembly, activation and lysine selection by the SIN3B histone deacetylase complex. Nature Communications 14, 2556 (2023).

18. Jamaladdin, S. et al. Histone deacetylase (HDAC) 1 and 2 are essential for accurate cell division and the pluripotency of embryonic stem cells. Proc Natl Acad Sci U S A 111, 9840–5 (2014).

19. Chandru, A., Bate, N., Vuister, G.W. & Cowley, S.M. Sin3A recruits Tet1 to the PAH1 domain via a highly conserved Sin3-Interaction Domain. Scientific reports 8, 14689 (2018).

20. Sidoli, S. et al. One minute analysis of 200 histone posttranslational modifications by direct injection mass spectrometry. Genome research 29, 978–987 (2019).

21. Baker, I.M., Smalley, J.P., Sabat, K.A., Hodgkinson, J.T. & Cowley, S.M. Comprehensive Transcriptomic Analysis of Novel Class I HDAC Proteolysis Targeting Chimeras (PROTACs). Biochemistry 62, 645–656 (2023).

22. Alexa, A. & Rahnenführer, J. Gene set enrichment analysis with topGO. Bioconductor Improv 27, 1–26 (2009).

23. Schwanhäusser, B. et al. Global quantification of mammalian gene expression control. Nature 473, 337–342 (2011).

24. Millard, C.J., Fairall, L., Ragan, T.J., Savva, C.G. & Schwabe, J.W. The topology of chromatin-binding domains in the NuRD deacetylase complex. Nucleic acids research 48, 12972–12982 (2020).

25. Cao, R. & Zhang, Y. SUZ12 is required for both the histone methyltransferase activity and the silencing function of the EED-EZH2 complex. Molecular cell 15, 57–67 (2004).

26. Kuzmichev, A., Nishioka, K., Erdjument-Bromage, H., Tempst, P. & Reinberg, D. Histone methyltransferase activity associated with a human multiprotein complex containing the Enhancer of Zeste protein. Genes & development 16, 2893–2905 (2002).

27. Laherty, C.D. et al. Histone deacetylases associated with the mSin3 corepressor mediate mad transcriptional repression. Cell 89, 349–56 (1997).

28. Zhang, Y., Iratni, R., Erdjument-Bromage, H., Tempst, P. & Reinberg, D. Histone deacetylases and SAP18, a novel polypeptide, are components of a human Sin3 complex. Cell 89, 357–364 (1997).

29. Zhang, Y., LeRoy, G., Seelig, H.P., Lane, W.S. & Reinberg, D. The dermatomyositis-specific autoantigen Mi2 is a component of a complex containing histone deacetylase and nucleosome remodeling activities. Cell 95, 279–89 (1998).

30. Huttlin, E.L. et al. Dual proteome-scale networks reveal cell-specific remodeling of the human interactome. Cell 184, 3022–3040. e28 (2021).

31. van den Berg, D.L., et al. An Oct4-centered protein interaction network in embryonic stem cells. Cell stem cell 6, 369–381 (2010).

32. Yang, J. et al. Genome-wide analysis reveals Sall4 to be a major regulator of pluripotency in murine-embryonic stem cells. Proceedings of the National Academy of Sciences 105, 19756–19761 (2008).

33. Williams, K. et al. TET1 and hydroxymethylcytosine in transcription and DNA methylation fidelity. Nature 473, 343–8 (2011).

34. Yang, X., Zhang, F. & Kudlow, J.E. Recruitment of O-GlcNAc transferase to promoters by corepressor mSin3A: coupling protein O-GlcNAcylation to transcriptional repression. Cell 110, 69–80 (2002).

35. Watson, P.J. et al. Insights into the activation mechanism of class I HDAC complexes by inositol phosphates. Nat Commun 7, 11262 (2016).

36. Kim, Y.J. et al. HDAC inhibitors induce transcriptional repression of high copy number genes in breast cancer through elongation blockade. Oncogene 32, 2828–35 (2013).

37. Leitch, H.G. et al. Naive pluripotency is associated with global DNA hypomethylation. Nature Structural & Molecular Biology 20, 311–316 (2013).

38. Abramson, J. et al. Accurate structure prediction of biomolecular interactions with AlphaFold 3. Nature, 1–3 (2024).

39. van Mierlo, G., Veenstra, G.J.C., Vermeulen, M. & Marks, H. The complexity of PRC2 subcomplexes. Trends in cell biology 29, 660–671 (2019).

40. Grant, P.A., Winston, F. & Berger, S.L. The biochemical and genetic discovery of the SAGA complex. Biochimica et Biophysica Acta (BBA)-Gene Regulatory Mechanisms 1864, 194669 (2021).

41. Jiang, H. The complex activities of the SET1/MLL complex core subunits in development and disease. Biochimica et Biophysica Acta (BBA)-Gene Regulatory Mechanisms 1863, 194560 (2020).

42. Banks, C.A. et al. Differential HDAC1/2 network analysis reveals a role for prefoldin/CCT in HDAC1/2 complex assembly. Scientific reports 8, 13712 (2018).

43. Bantscheff, M. et al. Chemoproteomics profiling of HDAC inhibitors reveals selective targeting of HDAC complexes. Nature biotechnology 29, 255–265 (2011).

44. Joshi, P. et al. The functional interactome landscape of the human histone deacetylase family. Molecular systems biology 9, 672 (2013).

45. Vcelkova, T. et al. GSE1 links the HDAC1/CoREST co-repressor complex to DNA damage. Nucleic Acids Research 51, 11748–11769 (2023).

46. Zhu, C. et al. Targeting the catalytic activity of HDAC1 in T cells protects against experimental autoimmune encephalomyelitis. bioRxiv, 2023.04.14.536700 (2023).

47. Hassig, C.A. et al. A role for histone deacetylase activity in HDAC1-mediated transcriptional repression. Proceedings of the National Academy of Sciences 95, 3519–3524 (1998).

48. Hess, L. et al. A toolbox for class I HDACs reveals isoform specific roles in gene regulation and protein acetylation. PLOS Genetics 18, e1010376 (2022).

49. Weerasinghe, S.V.W., Estiu, G., Wiest, O. & Pflum, M.K.H. Residues in the 11 Å Channel of Histone Deacetylase 1 Promote Catalytic Activity: Implications for Designing Isoform-Selective Histone Deacetylase Inhibitors. Journal of Medicinal Chemistry 51, 5542–5551 (2008).

50. Moser, M.A., Hagelkruys, A. & Seiser, C. Transcription and beyond: the role of mammalian class I lysine deacetylases. Chromosoma 123, 67–78 (2014).

51. Wang, S. et al. A potential histone-chaperone activity for the MIER1 histone deacetylase complex. Nucleic Acids Research 51, 6006–6019 (2023).

52. Dovey, O.M., Foster, C.T. & Cowley, S.M. Histone deacetylase 1 (HDAC1), but not HDAC2, controls embryonic stem cell differentiation. Proc Natl Acad Sci U S A 107, 8242–7 (2010).

